# Green autofluorescence intensity of skin and fingernails: A novel biomarker for non-invasive evaluation of pathological state of blood vessels

**DOI:** 10.1101/403832

**Authors:** Mingchao Zhang, Danhong Wu, Yue Tao, Yujia Li, Shufan Zhang, Xin Chen, Weihai Ying

**Affiliations:** Med-X Research Institute and School of Biomedical Engineering, Shanghai Jiao Tong University, Shanghai 200030, P.R. China; Department of Neurology, Shanghai Fifth People’s Hospital, Fudan University, Shanghai, P.R. China; Collaborative Innovation Center for Genetics and Development, Shanghai 200043, P.R. China

**Keywords:** Vascular health, Skin, Autofluorescence, Asymmetry, Skin, Diagnosis.

## Abstract

Stroke and myocardial infarction (MI) are two leading causes of death around the world. It is of great significance to establish novel and non-invasive approaches for evaluating pathological state of blood vessels, so that early interventions may be carried out to prevent incidence of stroke or MI. Our recent studies have suggested that altered ‘Pattern of Autofluorescence (AF)’ of skin and fingernails are novel biomarkers of acute ischemic stroke (AIS) and MI. In particular, our studies have shown characteristic increases in the green AF intensity of the fingernails and certain regions of the skin of AIS patients and MI patients. By determining the skin’s green AF of the Healthy Group, the Low-Risk Group for Developing AIS, and the High-Risk Group for Developing AIS, our current study has indicated that the green AF intensity in the right and left Ventroforefingers and Dorsal Antebrachium, the right Dorsal Centremetacarpus, Dorsal Index Fingers and Centremetacarpus, as well as the left Index Fingernails and Ventriantebrachium, is highly correlated with the risk of developing AIS. Our study has also suggested that oxidative stress and inflammation may account for the increased AF intensity in the Low-Risk Group and the High-Risk Group. These findings have suggested that the green AF intensity of the fingernails and certain regions of the skin is a novel biomarker for non-invasive evaluation of the injury levels of blood vessels and the risk of developing AIS or MI.

## Introduction

Non-invasive evaluation of pathological state of blood vessels and non-invasive prediction of the risk of developing acute ischemic stroke (AIS) and myocardial infarction (MI) is of great clinical significance. The evaluation and prediction is required for early intervention and diagnosis of AIS and MI. So far there is been no such an approach around the globe.

Our recent studies have suggested that characteristic ‘Pattern of Autofluorescence (AF)’ could be a novel biomarker for non-invasive diagnosis of AIS (3), MI (25), stable coronary artery disease (25), Parkinson’s disease (4) and lung cancer (14). In particular, there are marked and characteristic increases in the green AF intensity in the fingernails and certain regions of the skin of the AIS patients (3) and MI patients (25). Since oxidative stress can lead to increased epidermal green AF by altering keratin 1 (12, 13), the increased oxidative stress of AIS patients (2, 6, 23) and MI patients (1, 5, 11, 15, 24) may be responsible for the increased AF intensity. Our latest study has also indicated that inflammation can produced increased epidermal green AF intensity (26).

A number of studies have shown increased oxidative stress in the blood of the patients of diabetes (8, 18, 19) and hypertension (10, 16), which is an important pathogenic factor of diabetes (8), hypertension (10) and other cardiovascular diseases (10). Therefore, the persons with major risk factors for developing AIS or MI, including having such diseases as diabetes or hypertension, may also have the oxidative stress-induced increases in the green AF intensity at their fingernails or certain regions of their skin. It is also reasonable to propose that the more risk factors a person has, the higher oxidative stress in his body, which may lead to higher green AF intensity in their fingernails and/or certain regions of their skin. Based on this information, we propose our hypothesis that the green AF intensity in fingernails and certain regions of skin may become a novel biomarker for non-invasive evaluation of the injury levels of blood vessels as well as the risk to develop AIS or MI. We tested this hypothesis in our current study, which has provided evidence supporting this hypothesis.

## Methods and materials

### Human subjects

The study was conducted according to a protocol approved by the Ethics Committee of Shanghai Fifth People’s Hospital Affiliated to Fudan University. The human subjects in our study were divided into three groups: Group 1: The Healthy Group; Group 2: Low-Risk Group; and High-Risk Group 3. The following eight factors were used to determine if a person is in low-risk or high-risk of developing AIS: Diabetes, hypertension, atrial fibrillation or significant heart rate dissonance, obesity, smoking, abnormal triglyceride level in blood, family history of stroke, and lack of physical activity. The person with one or two of the factors is defined as ‘Low-Risk Person’, while the person with more than two of the factors is defined as ‘High-Risk Person’. The average age of Group 1, Group 2 and Group 3 is 65.66±1.16, 68.27± 0.58, and 68.69±0.72 years of old, respectively. The age of all of the subjects ranges from 50 - 80 years of old.

### Determinations of the Autofluorescence of Skin and Fingernails

A portable AF imaging equipment was used to detect the AF of the fingernails and certain regions of the skin of the human subjects. The excitation wavelength is 485 nm, and the emission wavelength is 500 - 550 nm. For all of the human subjects, the AF intensity in the following seven regions on both hands, i.e., fourteen regions in total, was determined, including the Index Fingernails, Ventroforefingers, Dorsal Index Finger, Centremetacarpus, Dorsal Centremetacarpus, Ventribrachium and Dorsal Antebrachium.

### Statistical analyses

All data are presented as mean ± SEM. Data were assessed by one-way ANOVA, followed by Student - Newman - Keuls *post hoc* test, except where noted. *P* values less than 0.05 were considered statistically significant.

## Results

### 1. Correlation between skin’s green AF intensity and risk of developing AIS

We determined the correlation between skin’s green AF intensity and risk of developing AIS. We found that the correlation between the AF intensity at right Ventroforefinger and the risk of developing AIS reaches 0.9924, while the correlation between the AF intensity at left Ventroforefinger and the risk of developing AIS is 0.9787 (Fig. 1). The correlation between the AF intensity at right and left Dorsal Antebrachium and the risk of developing AIS is 0.9825 and 0.9591, respectively (Fig. 2). While the correlation between the AF intensity at right Dorsal Centremetacarpus and the risk of developing AIS is 0.9971, the correlation between the AF intensity at left Dorsal Centremetacarpus and the risk of developing AIS is only 0.8241 (Fig. 3). The correlation between the AF intensity at right Dorsal Index Fingers and the risk of developing AIS reaches 0.949, while the correlation between the AF intensity at left Dorsal Index Fingers and the risk of developing AIS is only 0.8976 (Fig. 4). Similarly, the correlation between the AF intensity at right Centremetacarpus and the risk of developing AIS is 0.9734, while the correlation between the AF intensity at left Centremetacarpus and the risk of developing AIS is only 0.8867 (Fig. 5). We also found that the correlation between the AF intensity at right Ventriantebrachium and the risk of developing AIS is 0.7734, while the correlation between the AF intensity at left Ventriantebrachium and the risk of developing AIS is 0.9473 (Fig. 6).

**Fig 1.**
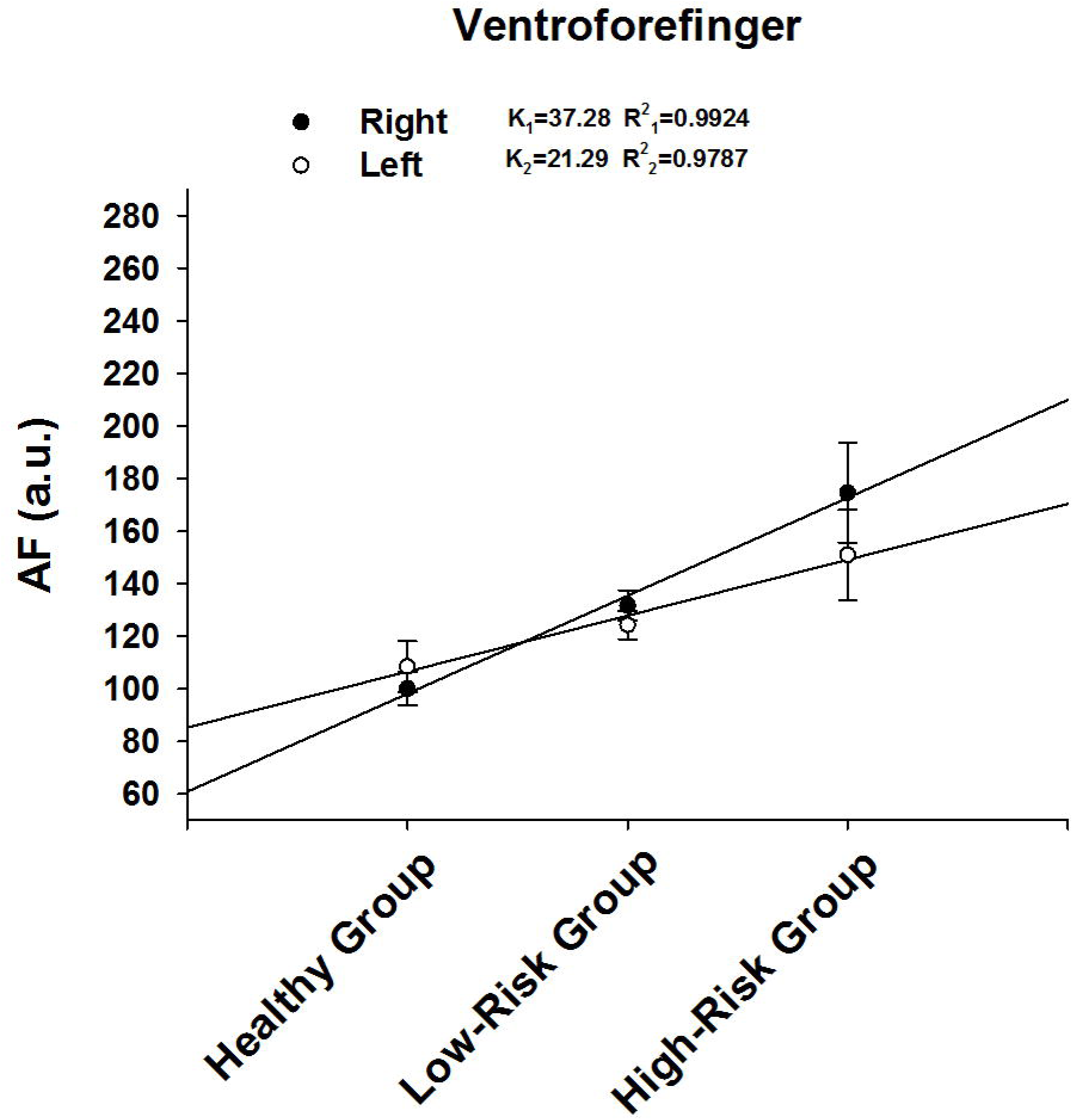
Correlation between the green AF intensity of Ventroforefinger’s skin and risk of developing AIS. The correlation between the AF intensity at right Ventroforefinger and the risk of developing AIS reaches 0.9924, while the correlation between the AF intensity at left Ventroforefinger and the risk of developing AIS is 0.9787. The number of subjects in the Healthy group, Low-Risk group and High-Risk group is 59, 236, and 109, respectively.

**Fig 2.**
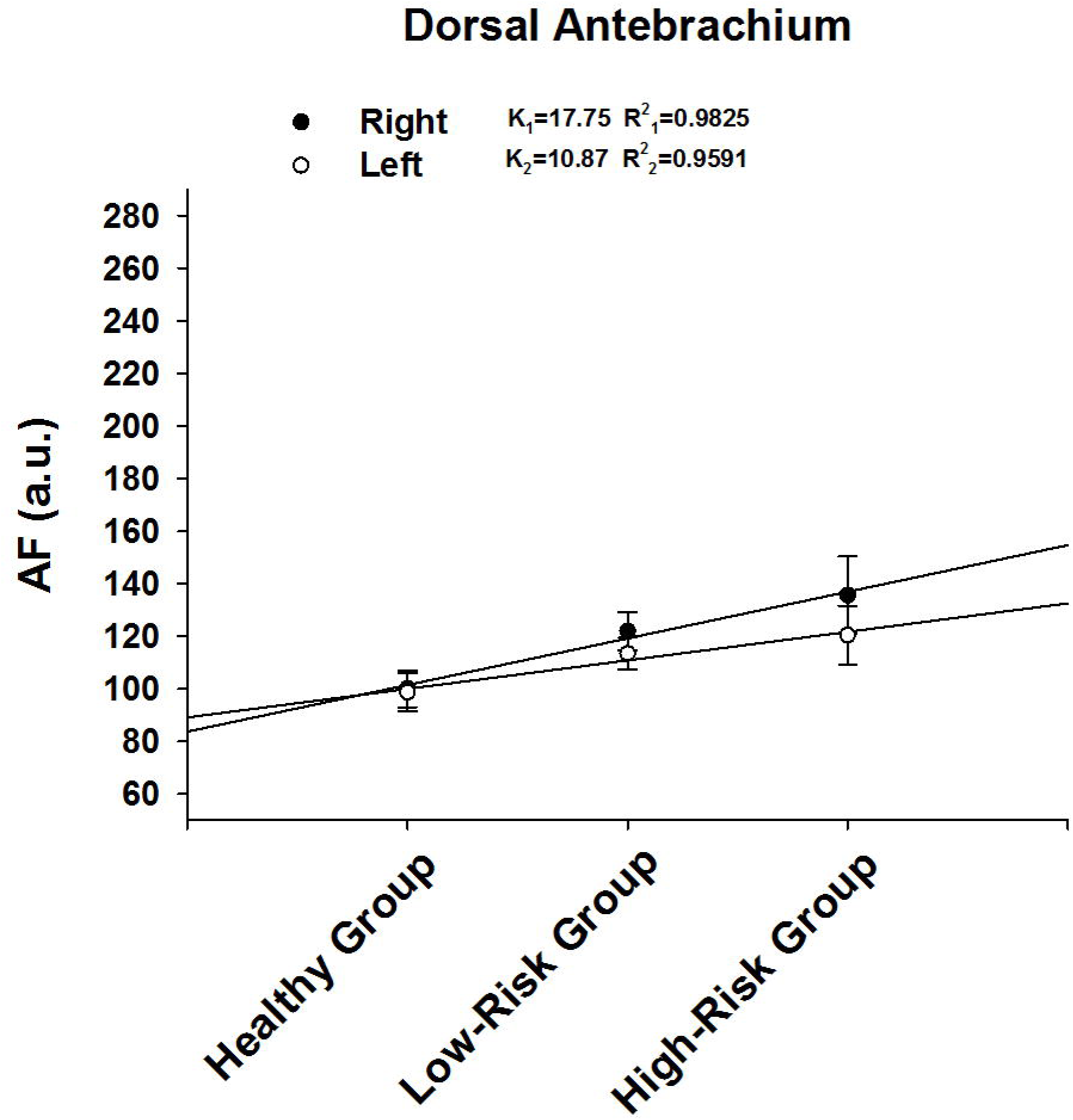
Correlation between the green AF intensity of Dorsal Antebrachium’ skin and risk of developing AIS. The correlation between the AF intensity at right and left Dorsal Antebrachium and the risk of developing AIS is 0.9825 and 0.9591, respectively. The number of subjects in the Healthy group, Low-Risk group and High-Risk group is 59, 236, and 109, respectively.

**Fig 3.**
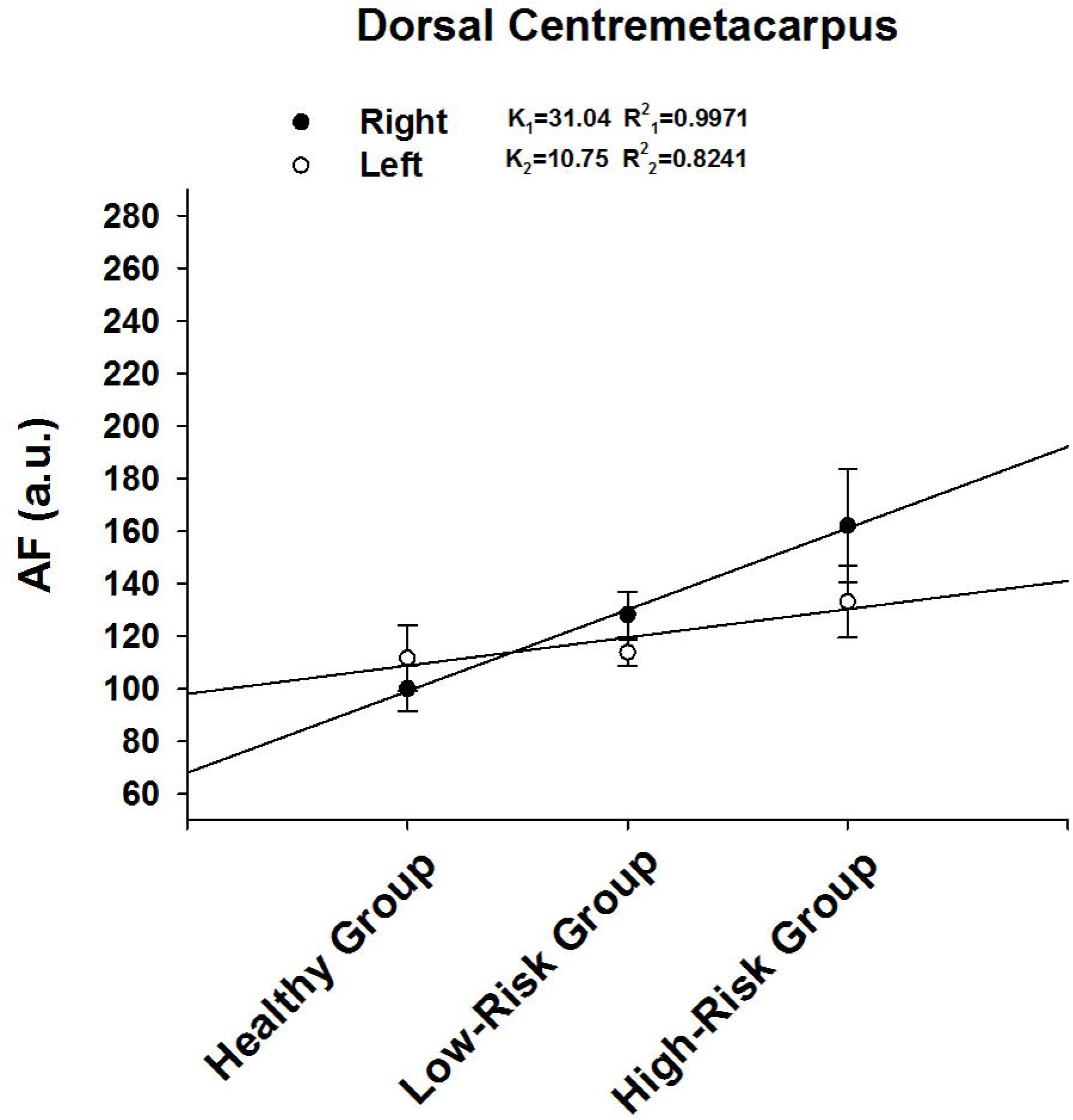
There is correlation only between the green AF intensity of right Dorsal Centremetacarpus’ skin and risk of developing AIS. While the correlation between the AF intensity at right Dorsal Centremetacarpus and the risk of developing AIS is 0.9971, the correlation between the AF intensity at left Dorsal Centremetacarpus and the risk of developing AIS is only 0.8241. The number of subjects in the Healthy group, Low-Risk group and High-Risk group is 59, 236, and 109, respectively.

**Fig 4.**
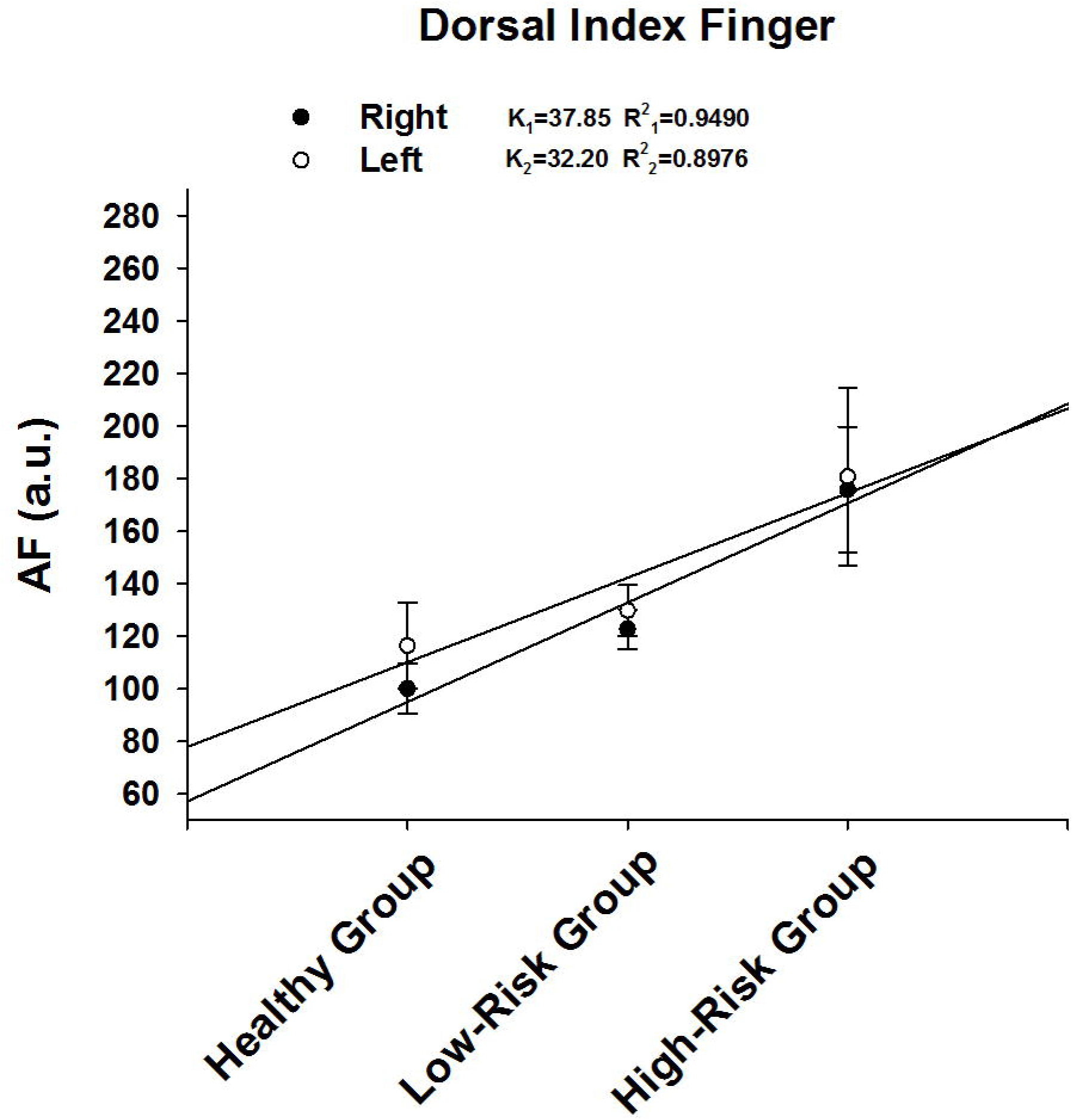
Correlation between the green AF intensity of Dorsal Index Fingers’ skin and risk of developing AIS. The correlation between the AF intensity at right Dorsal Index Fingers and the risk of developing AIS reaches 0.9490, while the correlation between the AF intensity at left Dorsal Index Fingers and the risk of developing AIS is only 0.8976. The number of subjects in the Healthy group, Low-Risk group and High-Risk group is 59, 236, and 109, respectively.

**Fig 5.**
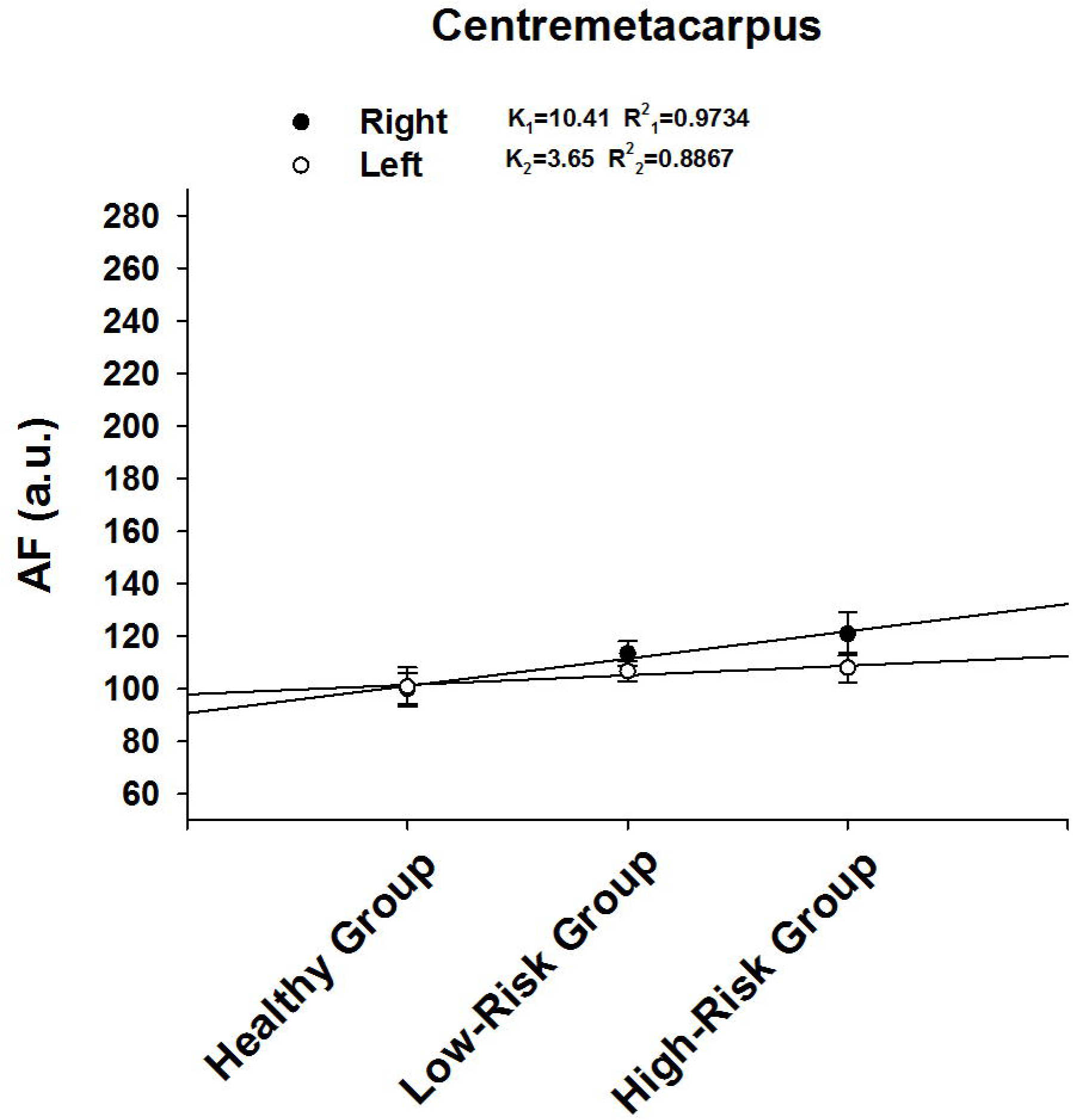
There is correlation between the green AF intensity of right Centremetacarpus’ skin and risk of developing AIS. The correlation between the AF intensity at right Centremetacarpus and the risk of developing AIS is 0.9734, while the correlation between the AF intensity at left Centremetacarpus and the risk of developing AIS is 0.8867. The number of subjects in the Healthy group, Low-Risk group and High-Risk group is 59, 236, and 109, respectively.

**Fig 6.**
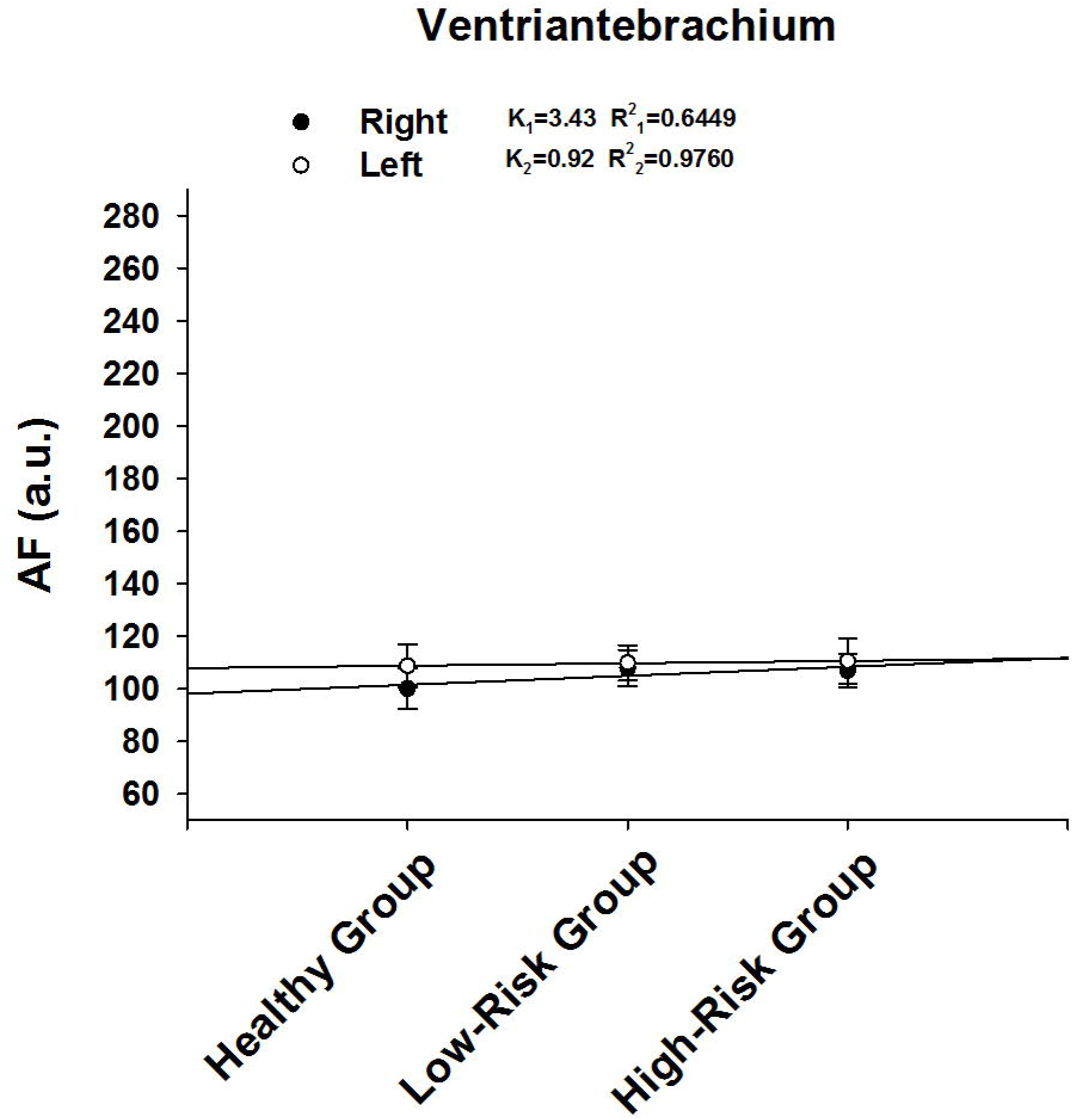
There is correlation between the green AF intensity of left Ventriantebrachium’ skin and risk of developing AIS. The correlation between the AF intensity at right Ventriantebrachium and the risk of developing AIS is 0.7734, while the correlation between the AF intensity at left Ventriantebrachium and the risk of developing AIS is 0.9473. The number of subjects in the Healthy group, Low-Risk group and High-Risk group is 59, 236, and 109, respectively.

### 2. Correlation between the green AF intensity of the left Index Fingernails and risk of developing AIS

We determined the correlation between Fingernails’ green AF intensity and risk of developing AIS. We found that the correlation between the AF intensity at right Index Fingernails and the risk of developing AIS is 0.7734, and the correlation between the AF intensity at left Index Fingernails and the risk of developing AIS also reaches 0.9473 (Fig. 7).

**Fig 7.**
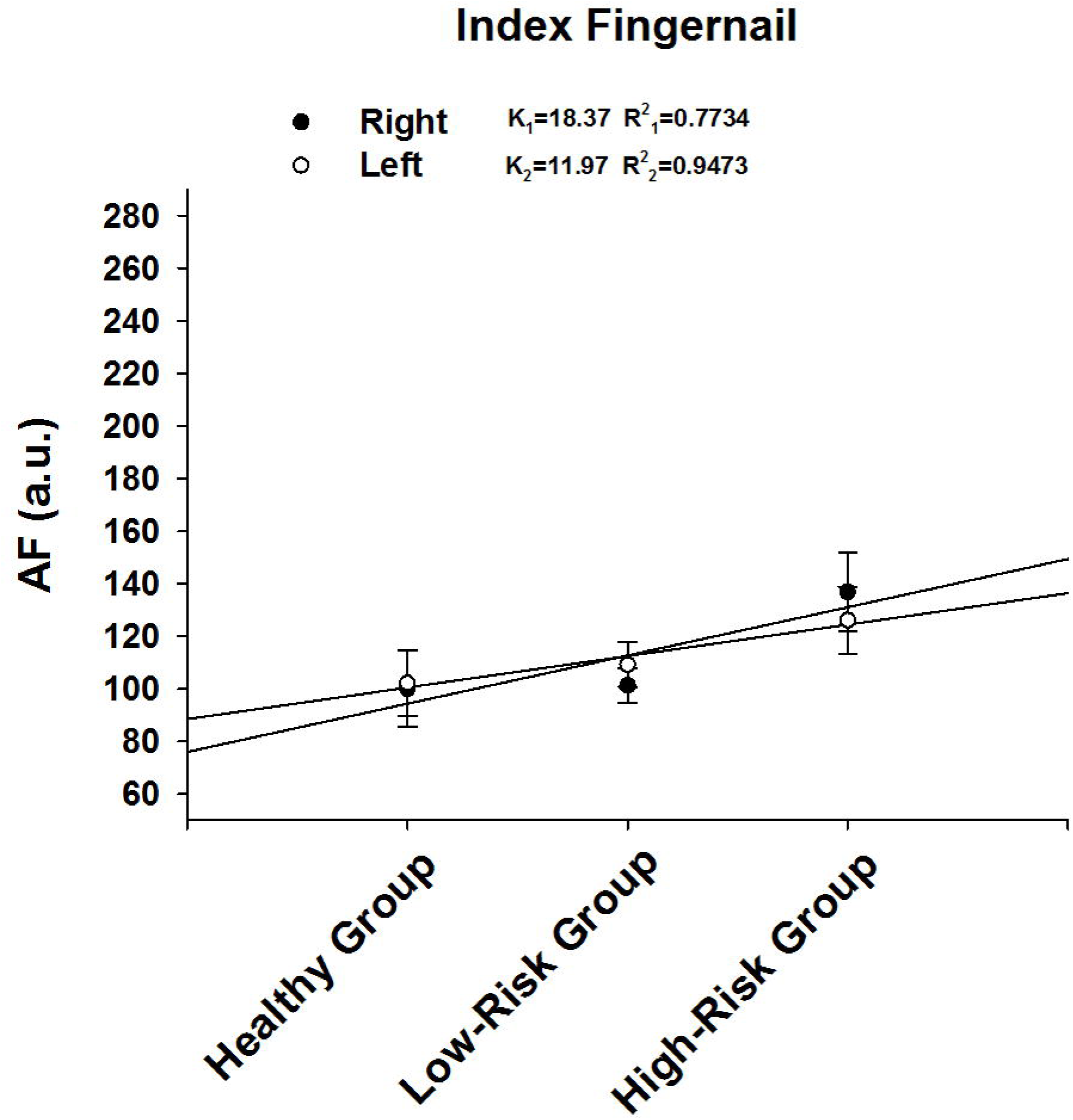
There is correlation between the green AF intensity of left Index Fingernails and risk of developing AIS. The correlation between the AF intensity at right Index Fingernails and the risk of developing AIS reaches 0.7734, and the correlation between the AF intensity at left Index Fingernails and the risk of developing AIS also reaches 0.9473. The number of subjects in the Healthy group, Low-Risk group and High-Risk group is 59, 236, and 109, respectively.

## Discussion

The major findings of our study include: The green AF intensity of fingernails and certain regions of skin is highly correlated with the risk of developing AIS. These regions include the right and left Ventroforefingers and Dorsal Antebrachium, the right Dorsal Centremetacarpus, Dorsal Index Fingers and Centremetacarpus, as well as the left Index Fingernails and Ventriantebrachium.

Non-invasive evaluation of injury levels of blood vessels and prediction of the risk of developing AIS or MI is of great clinical significance. This evaluation and prediction is required for early intervention and prevention of these diseases. Our current study has suggested that the epidermal green AF intensity of the fingernails and certain regions of skin may become a novel biomarker for non-invasive evaluation of both the injury levels of blood vessels and risk of developing AIS.

Our previous studies have indicated that the increased epidermal green AF indicates the injury levels produced by oxidative stress or inflammation, instead merely a probability of injury: 1) We have found that in healthy persons (3) and normal mice that are not exposed to UV (12, 13) or inflammation-inducing agents (26), the basal epidermal green AF is at very low levels. 2) Such insults as UV (12, 13) and LPS (26) can dose-dependently increase the epidermal AF intensity. 3) Compared with Low-Risk persons – the persons with relatively mild cardiovascular diseases such as hypertension, severe cardiovascular diseases and cerebrovascular diseases such as AIS can produce significant increases in the AF intensity (3). 4) We have found that keratin 1 – a major cytoskeletal protein – is rapidly degraded by such oxidative stress generators as UV (12, 13). Therefore, our green AF-based evaluation method is not merely a method of ‘risk evaluation’. Instead, our green AF-based method is essentially an ‘damage-detection method’, which has surpassed previous methods of ‘risk evaluation’.

Our current study has been conducted to determine the correlation between the AF intensity of each region examined and the risk of developing AIS. Our study has shown strong correlation between the risk of developing AIS and the green AF intensity of fingernails and certain regions of skin. These findings have suggested that the green AF intensity at fingernails and certain regions of the skin can become not only an ‘Index of Injury’, but also a novel biomarker for non-invasive evaluation of the risk of developing AIS.

A key scientific question for our current finding is: What is the mechanism underlying the increases in the AF intensity in the Low-Risk Group and the High-Risk Group? Based on previous findings from our lab and other groups, it appears that increased oxidative stress in the body of the patients of these two groups at least partially mediates the AF increases: First, our previous study has suggested that oxidative stress induced by such factors as UV can lead to increased epidermal green autofluorescence by altering keratin 1 (12, 13). Second, a number of studies have shown increased oxidative stress in the blood of the patients of diabetes (8, 18, 19) and hypertension (10, 16). Third, multiple studies have also suggested that oxidative stress is an important pathogenic factor of diabetes (8), hypertension (10) and other cardiovascular diseases (10). Fourth, there is also evidence indicating positive correlation between the levels of oxidative stress and the severity of such diseases as diabetes (22, 27). Therefore, it is reasonable to propose that the persons with more risk factors to develop AIS or MI may have higher levels of oxidative stress in their body and/or longer durations of increased oxidative stress in their body.

There is evidence suggesting that the increased levels of inflammation in the body of the persons of the Low-Risk Group and the High-Risk Group may also be causative to the increased green AF intensity: 1) Our latest study has indicated that inflammation can also produce increased epidermal green AF intensity (26); and 2) previous studies have indicated that the patients of diabetes (7, 9, 21) and hypertension (17, 20) have increased inflammation levels.

Collectively, these findings have suggested that our AF intensity-based method could be used to provide non-invasive and rapid evaluation of the extent of injury of a person’s blood vessels and a person’s risk of developing AIS or MI. Since our AF imaging device is portable and inexpensive, the device may become the first non-invasive and portable device for evaluating the extent of injury of a person’s blood vessels as well as a person’s risk of developing AIS or MI. It is expected that with the applications of big data science and artificial intelligence (AI), as well as the increases in the regions of skin examined for AF, increasingly higher precision in the evaluation may be achieved in the future.

## Acknowledgment

The authors would like to acknowledge the financial support by a Major Special Program Grant of Shanghai Municipality (Grant # 2017SHZDZX01) (to W.Y.) and a Major Research Grant from the Scientific Committee of Shanghai Municipality #16JC1400502 (to W.Y.).

## References

1. Caimi G, Canino B, Incalcaterra E, Ferrera E, Montana M, Lo Presti R. Behaviour of protein carbonyl groups in juvenile myocardial infarction. Clin Hemorheol Microcirc 53: 297–302, 2013.

2. Ciancarelli I, Di Massimo C, De Amicis D, Carolei A, Tozzi Ciancarelli MG. Evidence of redox unbalance in post-acute ischemic stroke patients. Curr Neurovasc Res 9: 85–90, 2012.

3. Wu D, Zhang M, Tao Y, Li Y, Zhang S, Chen X, Ying W. Asymmetric increases in the intensity of the green autofluorescence of ischemic stroke patients’ skin and fingernails: A novel diagnostic biomarker for ischemic stroke. bioRxiv 310904, 2018.

4. Wu D, Zhang M, Tao Y, Li Y, Shen L, Li Y, Ying W. Selectively increased autofluorescence at fingernails and certain regions of skin: A potential novel diagnostic biomarker for Parkinson disease. bioRxiv 322222, 2018.

5. Deepa M, Pasupathi P, Sankar KB, Rani P, Kumar SP. Free radicals and antioxidant status in acute myocardial infarction patients with and without diabetes mellitus. Bangladesh Med Res Counc Bull 35: 95–100, 2009.

6. Dominguez C, Delgado P, Vilches A, Martin-Gallan P, Ribo M, Santamarina E, Molina C, Corbeto N, Rodriguez-Sureda V, Rosell A, Alvarez-Sabin J, Montaner J. Oxidative stress after thrombolysis-induced reperfusion in human stroke. Stroke 41: 653–60, 2010.

7. Duncan BB, Schmidt MI. Chronic activation of the innate immune system may 12 underlie the metabolic syndrome. Sao Paulo Med J 119: 122–7, 2001.

8. Evans JL, Goldfine ID, Maddux BA, Grodsky GM. Oxidative stress and stress-activated signaling pathways: a unifying hypothesis of type 2 diabetes. Endocr Rev 23: 599–622, 2002.

9. Grimble RF. Inflammatory status and insulin resistance. Curr Opin Clin Nutr Metab Care 5: 551–9, 2002.

10. Houston MC. Nutrition and nutraceutical supplements in the treatment of hypertension. Expert Rev Cardiovasc Ther 8: 821–33, 2010.

11. Madole MB, Bachewar NP, Aiyar CM. Study of oxidants and antioxidants in patients of acute myocardial infarction. Adv Biomed Res 4: 241, 2015.

12. Zhang M, Dhruba TM, He H, Li Y, Yan W, Yan W, Zhu Y, Ying W. UV-Induced Keratin 1 Proteolysis Mediates UV-Induced Skin Damage. bioRxiv 226308, 2018.

13. Zhang M, Dhruba TM, Li Y, Ying W. Oxidative stress mediates UVC-induced increases in epidermal autofluorescence of C57 mouse ears. BioRxiv 298000, 2018.

14. Zhang M, Tao Y, Chang Q, Li Y, Chu T, Ying W. Selectively increased autofluorescence at certain regions of skin may become a novel diagnostic biomarker for lung cancer. bioRxiv 315440, 2018.

15. Mocatta TJ, Pilbrow AP, Cameron VA, Senthilmohan R, Frampton CM, Richards AM, Winterbourn CC. Plasma concentrations of myeloperoxidase predict mortality after myocardial infarction. J Am Coll Cardiol 49: 1993–2000, 2007.

16. Montezano AC, Dulak-Lis M, Tsiropoulou S, Harvey A, Briones AM, Touyz RM. Oxidative stress and human hypertension: vascular mechanisms, biomarkers, and novel therapies. Can J Cardiol 31: 631–41, 2015.

17. Pietri P, Vlachopoulos C, Tousoulis D. Inflammation and Arterial Hypertension: From Pathophysiological Links to Risk Prediction. Curr Med Chem 22: 2754–61, 2015.

18. Rosen P, Nawroth PP, King G, Moller W, Tritschler HJ, Packer L. The role of oxidative stress in the onset and progression of diabetes and its complications: a summary of a Congress Series sponsored by UNESCO-MCBN, the American Diabetes Association and the German Diabetes Society. Diabetes Metab Res Rev 17: 189–212, 2001.

19. Samadi A, Gurlek A, Sendur SN, Karahan S, Akbiyik F, Lay I. Oxysterol species: reliable markers of oxidative stress in diabetes mellitus. J Endocrinol Invest, 2018.

20. Satou R, Penrose H, Navar LG. Inflammation as a Regulator of the Renin-Angiotensin System and Blood Pressure. Curr Hypertens Rep 20: 100, 2018.

21. Schmidt MI, Duncan BB, Sharrett AR, Lindberg G, Savage PJ, Offenbacher S, Azambuja MI, Tracy RP, Heiss G. Markers of inflammation and prediction of diabetes mellitus in adults (Atherosclerosis Risk in Communities study): a cohort study. Lancet 353: 1649–52, 1999.

22. Sedighi O, Makhlough A, Shokrzadeh M, Hoorshad S. Association between 14 plasma selenium and glutathione peroxidase levels and severity of diabetic nephropathy in patients with type two diabetes mellitus. Nephrourol Mon 6: e21355, 2014.

23. Seet RC, Lee CY, Chan BP, Sharma VK, Teoh HL, Venketasubramanian N, Lim EC, Chong WL, Looi WF, Huang SH, Ong BK, Halliwell B. Oxidative damage in ischemic stroke revealed using multiple biomarkers. Stroke 42: 2326–9, 2011.

24. Uppal N, Uppal V, Uppal P. Progression of Coronary Artery Disease (CAD) from Stable Angina (SA) Towards Myocardial Infarction (MI): Role of Oxidative Stress. J Clin Diagn Res 8: 40–3, 2014.

25. Qu X, Li Y, Tao Y, Zhang M, Wu D, Guan S, Han W, Ying W. Distinct Patterns of the Autofluorescence of Body Surface: Potential Novel Diagnostic Biomarkers for Stable Coronary Artery Disease and Myocardial Infarction. bioRxiv 330985, 2018.

26. Li Y, Zhang M, Tao Y, Ying W. Lipopolysaccharide (LPS) induces increased epidermal green autofluorescence of mouse. bioRxiv 501189, 2018.

27. Ziegler D, Sohr CG, Nourooz-Zadeh J. Oxidative stress and antioxidant defense in relation to the severity of diabetic polyneuropathy and cardiovascular autonomic neuropathy. Diabetes Care 27: 2178–83, 2004.

